# Intensive Chemotherapy Induces Cardiotoxicity via Reverse Electron Transport

**DOI:** 10.1101/2024.12.20.629711

**Authors:** Carla Pinto, Youzhong Liu, Sabrina Benaouadi, Yannis Sainte-Marie, Lotfi Laaouamir, Dimitri Marsal, Nick Van Gastel, Frédérique Savagner, Thomas Farge

## Abstract

Chemotherapy-induced cardiotoxicity has emerged as an important focus in oncology, driven by the growing number of cancer survivors. Intensive chemotherapies (iCT) used in the treatment of acute myeloid leukemia (AML) often lead to significant adverse cardiac events, which can reduce therapeutic benefits, limit treatment options, or even necessitate discontinuation—particularly for patients with pre-existing cardiac conditions, resulting in a loss of therapeutic opportunity. This study shows that the iCT triggers severe mitochondrial dysfunction in cardiac tissue, mirroring effects seen in ischemia-reperfusion models. Specifically, iCT results in succinate accumulation and elevated reactive oxygen species production, consistent with reverse electron transport phenomenon. Importantly, these effects were entirely prevented with RET inhibitors such as malonate or S1QEL1.1. *In vivo*, we demonstrate that malonate administration successfully prevents iCT-induced cardiotoxicity, maintaining left ventricular ejection fraction and fibrosis levels comparable to controls. Additionally, in an MLL-AF9-driven AML model, malonate sensitized leukemic cells to iCT. These findings support the dual potential of malonate: as an OXPHOS metabolism inhibitor to overcome chemoresistance in AML, while also reducing cardiotoxic risk for already vulnerable patients.

**One Sentence Summary:** This study demonstrates that malonate prevents chemotherapy-induced cardiotoxicity by inhibiting mitochondrial reverse electron transport (RET), preserving cardiac function, and simultaneously sensitizing AML cells to intensive chemotherapy.

## Introduction

Many cancer treatments are cardiotoxic, leading to cardiovascular diseases (CVD) that can cause significant mortality (*1–3*). Despite improved cancer survival rates, cardiotoxicity often surpasses cancer-related deaths (*4, 5*). Anthracyclines, such as doxorubicin, are well-known to induce reactive oxygen species (ROS) production and mitochondrial dysfunction, leading to cardiac cell death (*6*). However, most studies focus on anthracycline monotherapy, which does not reflect typical cancer treatment protocols, especially in acute myeloid leukemia (AML), where anthracyclines are used with cytarabine, another cardiotoxic chemotherapy (*1*). The incidence of cardiac events during intensive chemotherapy (iCT, Cytarabine + Doxorubicin) is particularly high in AML (*7, 8*).

Oxidative stress plays a central role in numerous pathologies, including heart failure (*9*). Antioxidant therapeutic strategies have proven highly effective in mitigating ischemia-reperfusion (I/R) damage as well as countering the cardiotoxic effects of antitumor therapies (*6, 10, 11*). This shared mechanism between these two models suggests that certain metabolic pathways are activated both by perfusion defects and by cardiotoxic agents. A key metabolic event in I/R induced heart failure is the accumulation of succinate, which drives pathological ROS production through the reverse electron transport (RET) within the mitochondrial electron transport chain (ETC) (*12*). This process involves succinate dehydrogenase (SDH) driving electrons from complex II (CII) back to complex I (CI), bypassing their usual flow to complex III (CIII). RET is well-documented in I/R injury, where it contributes to oxidative stress and tissue damage (*13*). Malonate, a competitive inhibitor of succinate dehydrogenase (SDH), effectively prevents succinate buildup and inhibits RET in multiple organs, including the heart (*14–19*).

Similarly to cardiac cells, AML blasts rely on oxidative metabolism for survival, especially under chemotherapy treatment (*20–25*). Interestingly, succinate has been described as an oncometabolite, crucial for cell survival and chemoresistance(*26, 27*). These shared pathological pathways suggest a dual benefit of targeting succinate metabolism: protecting cardiac function and enhancing chemotherapy efficacy.

## Results

### Doxorubicin and cytarabine combination drive major cardiotoxic effects

In the treatment of AML, patients follow the “3+7” regimen, which consists of three successive doses of anthracyclines (D1 to D3) and seven doses of AraC (D1 to D7). In this protocol, the toxic effects of iCT occur acutely in most cases (during chemotherapy induction)(*7*). Here, we used this regimen with doses of doxorubicin and AraC commonly employed in cardio-oncology studies (*28, 29*). We evaluated both chemotherapies alone or in combination in C57BL/6J mice (Fig. 1A). By day 7, the mice significantly lost weight when treated with iCT, unlike the mice in the monotherapy groups (Fig. 1B). An echocardiographic study allowed us to conclude the extreme toxicity of this model (Fig. 1C). When analyzed at a similar heart rate, the iCT group mice exhibited severe cardiac atrophy illustrated by a significant decrease in left ventricular volume and mass (Fig. 1D to E and Fig. S1A), as well as in cardiac output (Fig. 1F). Finally, we monitored the overall survival of the iCT-treated mice and observed a specific lethality in this group (Fig. 1G).

**Fig. 1.**
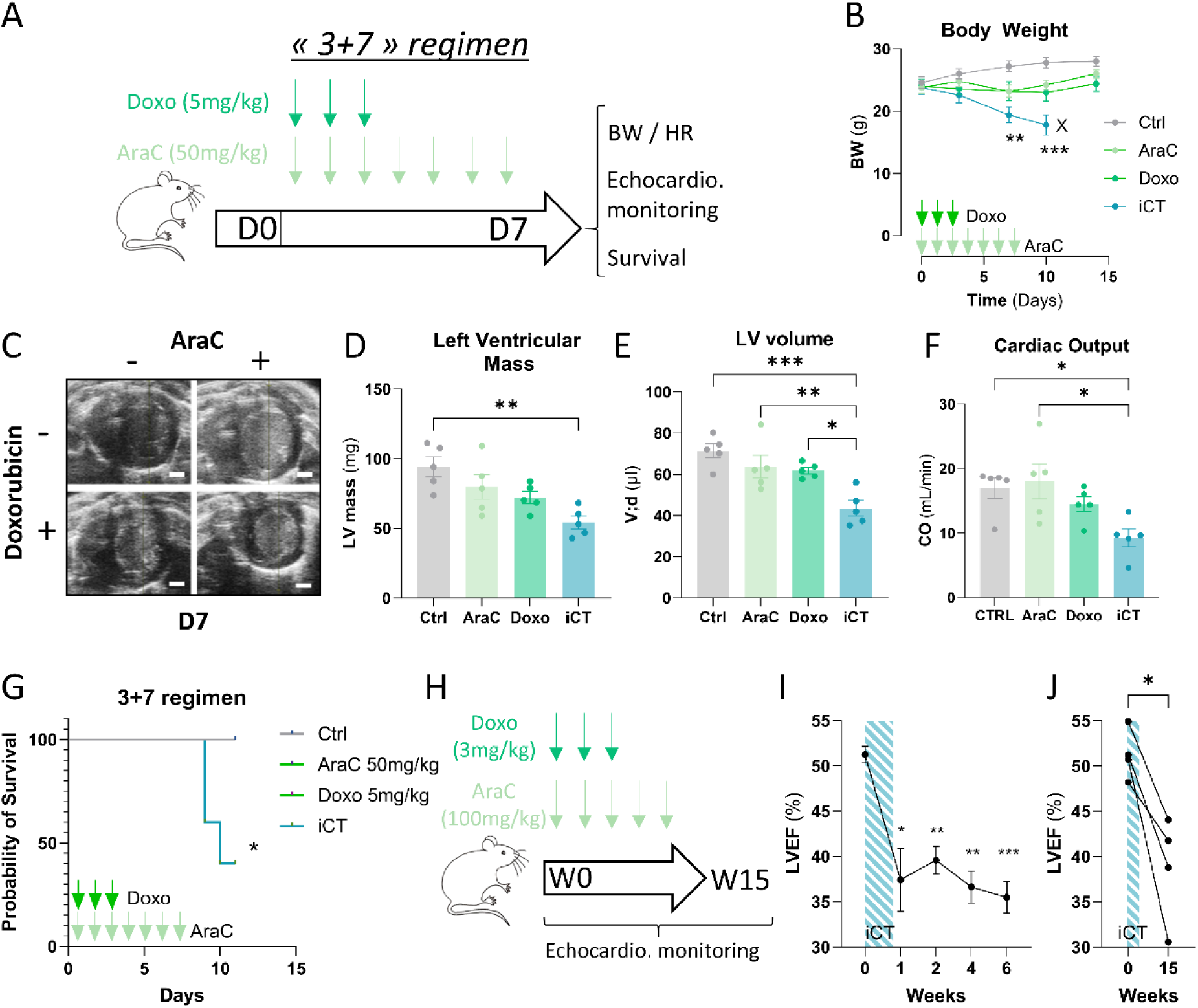
Intensive chemotherapies lead to higher cardiotoxicity compared to monotherapy treatments. (**A**) Experimental protocol for C57BL/6 mice treated with AraC, Doxo, or iCT following the 7+3 regimen and monitored using echocardiography. (**B**) Mouse Body Weight kinetics for 2 weeks treatment with AraC, Doxo, and iCT (n=5 mice per group). One-way ANOVA: **p<0,01; ***p<0,001. (**C**) Representative echocardiography images of mice treated with AraC, Doxo or iCT after 7 days. (**D**)Quantification of left ventricular volume after 7 days of treatment based on echocardiography (n=5 mice per group). One-way ANOVA: *p<0,05; **p<0,01; ***p<0,001. (**E**) Quantification of left ventricular mass on 7 days treated mice based on echocardiography (n=5 mice). One-way ANOVA: **p<0,01. (**F**) Cardiac output of 7 days treated mice based on echocardiography (n=5 mice). One-way ANOVA: *p<0,05. (**G**) Overall survival of AraC, Doxo and iCT treated mice (n=5mice). Log-rank test: *p<0,05. (**H**) Experimental protocol of C57Bl/6 mice treated with AraC, Doxo or iCT and monitored by echocardiography during 42 days. (**I**) (**K**) Kinetics of Left Ventricular Ejection Fraction in iCT-treated mice measured by echocardiography (n=6 mice per group). One-way ANOVA: *p<0,05; **p<0,01; ***p<0,001. (**J**) Kinetics of Left Ventricular Ejection Fraction in iCT-treated mice measured by echocardiography (n=6 mice per group). One-way ANOVA: *p<0,05.

In parallel, we conducted *in vitro* investigations to identify suitable conditions for studying cardiotoxicity mechanisms. We selected a 0.4 µM doxorubicin for 24 hours as optimal for H9C2 cardiomyoblastic cells, allowing metabolic adaptation without the massive cell death observed with the commonly used 1 µM dose (Fig. S1B). Transcriptomic data from *in vivo* doxorubicin-treated models reveal enrichment in mitochondrial oxidative metabolism signatures without glycolytic shifts, as observed in pig left ventricle (GSE197049) (*30, 31*) (Fig. S1C). At 0.4 µM, doxorubicin increased spare capacity without altering basal mitochondrial respiration, while the 1 µM dose significantly reduced it, aligning with prior studies (*32*) (Fig. S1D and E). Although 1 µM is commonly used in cardio-oncology, it induces major cell death and is rarely used in oncology (*33*). Our adapted AML reference chemotherapy combination (iCT) demonstrated greater toxicity than monotherapies, as shown *in vivo* (Fig. S1F).

The lethality observed with the “3+7” regimen, forced us to adjust the chemotherapy concentrations to those used in reference preclinical AML studies (*24*) (Fig. 1H). We thus established an iCT treatment model showing only a temporary reduction in body weight (Fig. S1G), accompanied by a decrease in left ventricular ejection fraction (LVEF) 7 days after treatment, which remains low even after 6 weeks (Fig. 1I). Finally, these same mice exhibited a reduced LVEF after 15 weeks (Fig. 1J), demonstrating that this model represents both acute and chronic cardiotoxicity. In conclusion, the combination of anthracycline with AraC results in increased toxicity to cardiac tissue in tested concentrations. Identifying the metabolic mechanisms involved in the cardiotoxicity of combined chemotherapies is essential for developing cardioprotective measures.

### Succinate is a key player in iCT-induced cardiotoxicity

To metabolically characterize the iCT cardiotoxicity model, we analyzed cardiac tissue using mass spectrometry to obtain its metabolome (Fig. 2A). Out of 105 detected metabolites, only 11 were significantly deregulated compared to PBS-treated mice, including the succinate/fumarate pair (Fig. 2B; Table 1). Specifically, succinate was significantly enriched while fumarate was depleted in these samples (Fig. 2C and D), indicating a metabolic effect of iCT on succinate dehydrogenase (SDH) as a source of iCT toxicity. Measuring protein expression of the catalytic subunit A of SDH reveals a significant overexpression of SDHA in heart post-iCT that could be related to increased SDH activity (Fig. 2E).

**Fig. 2.**
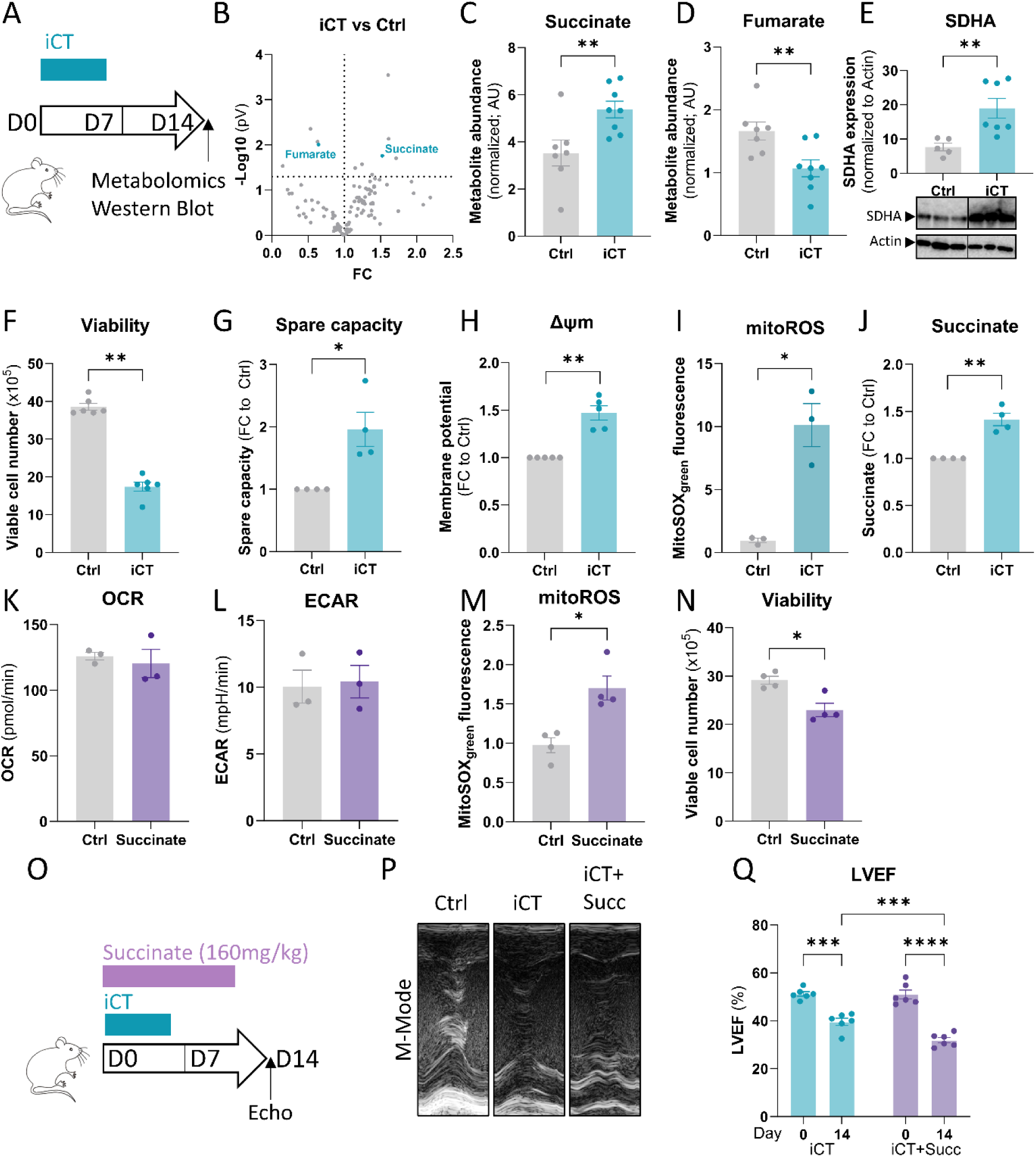
Metabolomic analysis linked succinate dehydrogenase activity to iCT-induced cardiotoxicity. (**A**) Experimental protocol for C57BL/6 mice treated with iCT following the 7+3 regimen. (**B**)Volcano plot of metabolite differences in mouse hearts between control and iCT-treated groups measured by mass spectrometry (n=7-8 mice). The x axis represents the Fold Change (FC) while the y axis represents –log10 (p-value <0,05). (**C**) Histogram of succinate metabolite levels abundance in iCT-treated Mouse Hearts (n=8 mice) compared to controls (n=7 mice). Mann-Whitney Test: **p<0,01. (**D**) Histogram of fumarate metabolite levels abundance in iCT-treated mouse hearts (n=8 mice) compared to controls (n=7 mice). Mann-Whitney Test: **p<0,01. (**E**) Western Blot analysis of SDHA expression normalized to actin expression in iCT-treated mouse hearts (n=7) compared to control hearts (n=5). Mann-Whitney Test: **p<0,01. (**F**) Histogram of viable cell counts in untreated versus iCT-treated H9C2 cells, measured using Malassez counting chamber (n=6). Mann-Whitney Test: **p<0,01. (**G**) Spare capacity measured by the Seahorse MitoStress Test in iCT-treated versus untreated H9C2 Cells (n=4). One-sample t-test: *p<0.05. (**H**) Histogram of mitochondrial membrane potential (Mitotracker red) in H9C2 either untreated or treated with iCT, malonate, or the combination of iCT and malonate for 24 hours (n=5). One-sample t-test: *p<0.05. (**I**) Incucyte analysis of MitoSOX Green Staining in untreated and iCT-treated H9C2 cells during 24 hours normalized to the number of cells (n=9). One-way ANOVA: **p<0.01. (**J**) Histogram of succinate concentration in untreated and iCT-treated H9C2 cells during 24 hours normalized to control (n=3). One sample t-test: **p<0,01. (**K**) Oxygen Consumption Rate (OCR) measured by Seahorse MitoStress Test in Diethyl Succinate-Ttreated versus untreated H9C2 cells (n=3). Mann-Whitney Test ns: not significant. (**L**) Extracellular Acidification Rate (ECAR) measured by Seahorse MitoStress Test in Diethyl Succinate-treated versus untreated H9C2 cells (n=3). Mann-Whitney Test ns: non-significant. (**M**) Incucyte analysis of MitoSOX Green Staining in untreated and dimethyl succinate-treated H9C2 cells during 24 hours normalized to the number of cells (n=4). Mann-Whitney Test: *p<0,05. (**N**) Histogram of viable cell counts in untreated versus diethyl succinate-treated H9C2 cells, measured using Malassez counting chamber (n=4). Mann-Whitney Test: *p<0,05. (**O**) Experimental protocol of C57Bl/6 mice treated with iCT and diethyl succinate for 7 and 14 days respectively, and monitored by echocardiography. (**P**) Representative M-mode echocardiographic images of mouse hearts, either untreated or treated with iCT and diethyl succinate for 7 or 14 days. (**Q**) Kinetics of Left Ventricular Ejection Fraction in mice treated with iCT or with both iCT and diethyl succinate, measured by echocardiography before and 14 days after treatment (n=6 mice per group). Two-way ANOVA: ***p<0,001; ****p<0,0001.

To further elucidate the underlying mechanisms observed *in vivo*, we characterized the impact of iCT using the rat cardiomyoblast H9C2 model *in vitro*. iCT reduced the viability of this cell line (Fig. 2F), accompanied by an increase in mitochondrial spare capacity, membrane potential and ROS (Fig. 2G to I). Intracellular succinate level was also increased upon iCT treatment (Fig. 2J). To assess the impact of this increased succinate concentration, we supplemented H9C2 culture medium with 5mM diethyl succinate (capable of entering cells passively). Seahorse analysis of these cells after 24 hours of culture showed no change in oxygen consumption rate compared to untreated cells (Fig. 2K and L). Succinate supplementation resulted in an increased production of mitochondrial ROS (Fig. 2M) and decreased cell viability (Fig. 2N).

Then, to understand the role of succinate in iCT cardiotoxicity *in vivo*, we treated mice with the chemotherapy combination as previously and added diethyl succinate treatment (160mg/kg, 10 doses over two weeks) for one group (Fig. 2O). Echocardiographic analysis revealed that succinate addition to iCT exacerbated LVEF reduction two weeks post-iCT (Fig. 2P and Q). Collectively, these results highlight the role of succinate accumulation as a driver of iCT induced heart failure, as a reminiscence of the reverse electron transport mechanisms observed in I/R models (*12, 34*).

### Malonate reverses iCT toxicity induced by succinate accumulation

The accumulation of succinate, coupled with the overexpression of SDHA, highlights complex II of the mitochondrial respiratory chain as a prime target for countering iCT-induced cardiotoxicity. Consequently, we used malonate, a competitive inhibitor of SDHA to block this pathological mechanism crucial in ischemia-induced heart failure but not yet observed under chemotherapeutic treatments (*14*). We used a relatively low dose of malonic acid (100 µM) in combination with iCT to counteract its toxicity on our *in vitro* model. After 24 hours of treatment, we demonstrated a protective effect of malonate, inhibiting cell death (Fig. 3A and B).

**Fig. 3.**
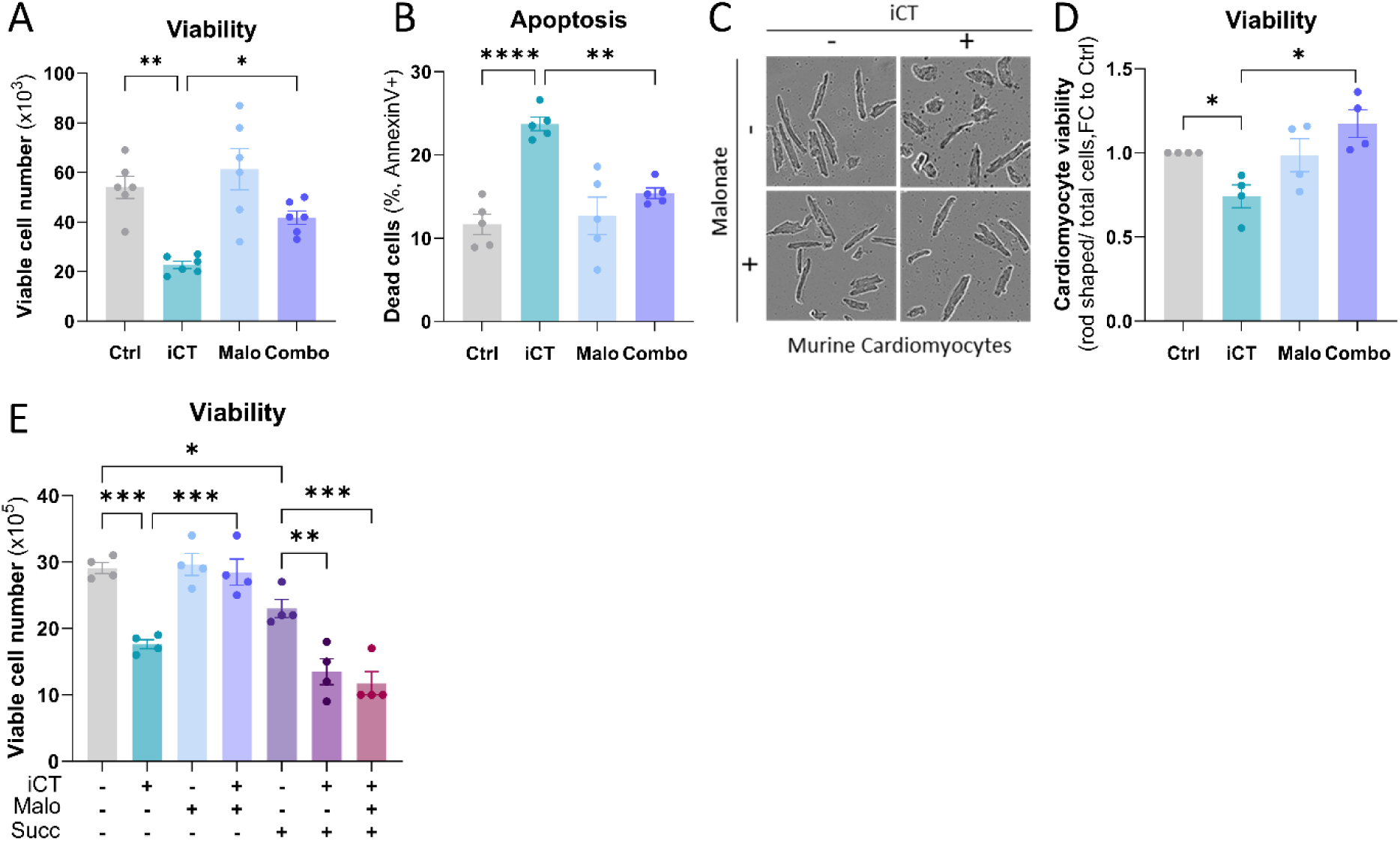
Malonate, an inhibitor of succinate dehydrogenase, counteracts iCT toxicity. (**A**) Histogram showing viable cell counts of H9C2 cells, either untreated or treated with iCT, malonate, or the combination of iCT and malonate (Combo), normalized to the control cell count, measured using a Malassez counting chamber (n=6). One-way ANOVA: *p<0.05; **p<0,01. (**B**) Flow cytometry analysis displaying the percentage of dead cells, marked as Annexin V^+^, in H9C2 cells either untreated or treated with iCT, malonate, or the combination of iCT and malonate (Combo) for 24 hours (n=5). One-way ANOVA: **p<0,01; ****p<0,0001. (**C**) Representative images of isolated murine cardiomyocytes, either untreated or treated with iCT, malonate, or a combination of both. (**D**) Histogram of cardiomyocyte viability, representing the percentage of rod-shaped cells among the total cell count, for isolated murine cardiomyocytes that were untreated or treated with iCT, malonate, or their combination for 24 hours (n=4) One sample t-test, *p<0,05. Histogram showing viable cell counts of H9C2 cells, either untreated or treated with iCT, malonate, diethyl succinate, or their combinations, measured using a Malassez counting chamber (n=4). One-way ANOVA: *p<0,05, **p<0,01, ***p<0,001.

Further, we treated murine isolated cardiomyocytes with iCT or malonate for 24 hours and observed similar results (Fig. 3C and D), demonstrating the cardioprotective capacity of malonate against iCT-induced cytotoxicity. Interestingly, malonate failed to counteract the toxicity of AraC or doxorubicin alone, underscoring the importance of focusing on clinically relevant combinations (Fig. S2A). We observed the same protective effect of dimethyl-malonate on H9C2 at different concentrations, especially with a 24-hour pre-treatment before adding iCT (Fig. S2B).

Additionally, we confirmed the toxicity of iCT and succinate alone or in combination while succinate addition reverses the protective effect of malonate (Fig. 3E). This demonstrates that the reactions catalyzed by SDHA are indeed the source of iCT toxicity.

### iCT increases ROS production through mitochondrial reverse electron transport

To specify the metabolic mechanisms at play following iCT and malonate treatments, we conducted series of experiments to assess mitochondrial functions. The analysis of basal and maximal mitochondrial respiration, as well as proton leak, ATP production and NAD^+^ levels demonstrated an increase in oxidative metabolism following iCT treatment. The addition of malonate led to a restored mitochondrial activity to control levels (Fig. 4A to D and S3A). Surprisingly, we also observed an accumulation of NADH levels following iCT treatment which was reversed by malonate (Fig. 4E). Additionally, iCT induced succinate accumulation, whereas malonate effectively maintained its level comparable to controls (Fig. 4F). The immediate increase in mitochondrial ROS production induced by iCT treatment and its reversal upon malonate treatment demonstrate that targeting SDH decrease mitochondrial oxidative stress (Fig. 4G and H). The results demonstrate an increased oxidative status following iCT treatment, which is reversed by SDH inhibition with malonate. This heightened metabolic activity may be linked to an increased NADH/NAD+ cycling, potentially saturating complex III and promoting RET.

**Fig. 4.**
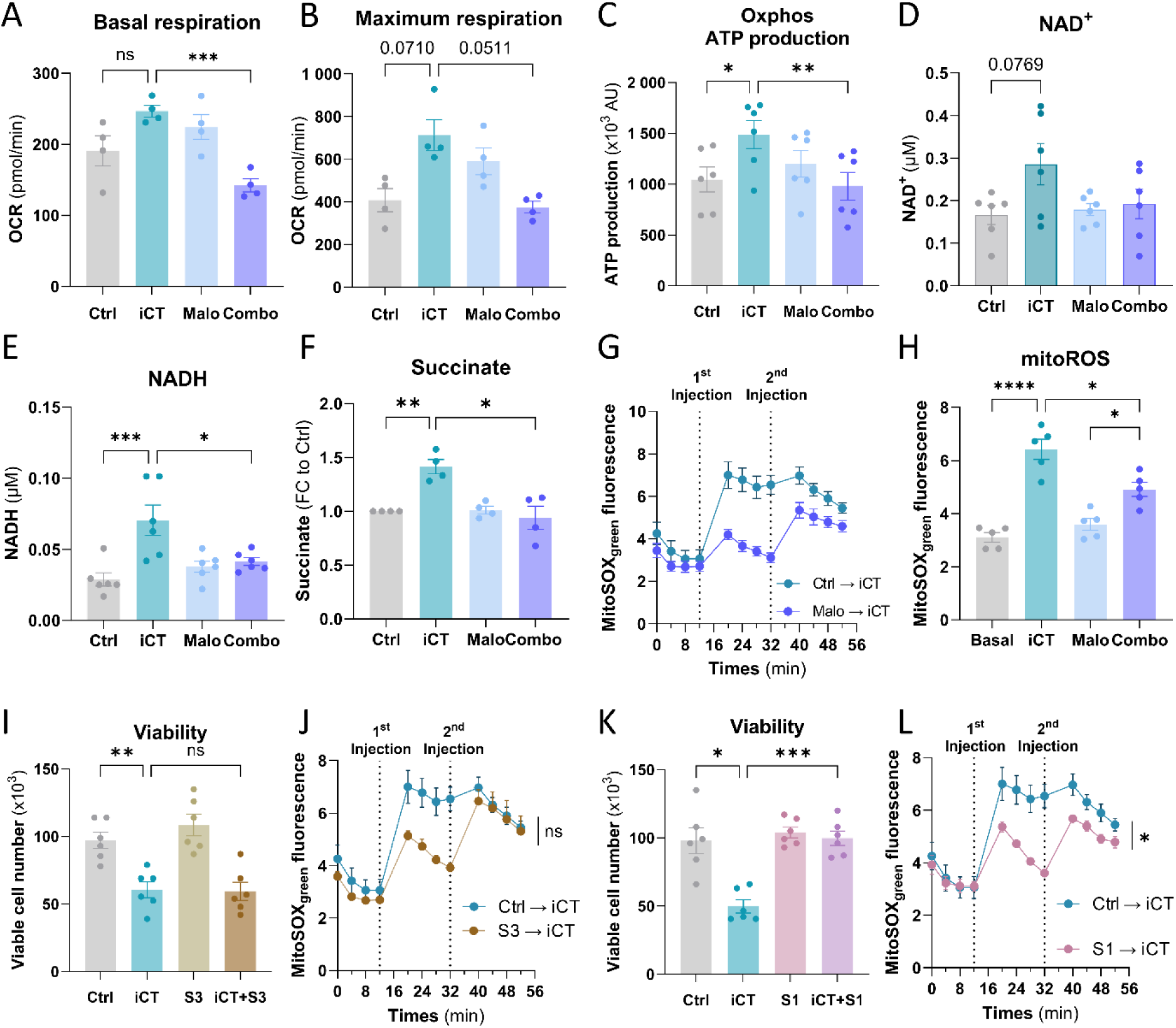
Malonate prevents iCT toxicity by disrupting reverse electron transport. (**A**) Basal oxygen consumption rate measured by Seahorse MitoStress Test in untreated H9C2 cells or those treated with iCT, malonate, or their combination (n=4) Brown-Forsythe and Welch ANOVA test: ***p<0,001. (**B**) Maximum oxygen consumption rate measured by Seahorse MitoStress Test in H9C2 either untreated or treated with iCT, malonate, or the combination of iCT and malonate (Combo) for 24 hours (n=4) Brown-Forsythe and Welch ANOVA test. (**C**) ATP production measured by Seahorse MitoStress Test in H9C2 cells either untreated or treated with iCT, malonate, or the combination of iCT and malonate (Combo) for 24 hours (n=6). Friedman Test, *p<0,05; **p<0,01. (**D**) Histogram of NAD^+^ concentration in H9C2 cells, either untreated or treated with iCT, malonate, or their combination (n=6). One-way ANOVA. (**E**) Histogram of NADH concentration in H9C2 cells, either untreated or treated with iCT, malonate, or their combination (n=6). One-way ANOVA: **p<0,01. (**F**) Histogram of succinate concentration in H9C2 cells treated with iCT, Malonate, or the combination of Malonate and iCT, normalized to control cells (n=4). One sample t-test, *p<0,05; **p<0.01. (**G**) Kinetics and (**H**) Histogram quantification of Mitosox Green fluorescence representing mitochondrial ROS before and after pre-treating H9C2 cells with malonate, followed by iCT treatment, or after iCT treatment alone (n=6). One-way ANOVA: *p<0.05; **p<0.01 (**I**) Histogram of viable H9C2 cell counts, comparing untreated cells to those treated with iCT, S3QEL2, or a combination of iCT and S3QEL2, measured with a Malassez counting chamber (n=6). Brown-Forsythe and Welch ANOVA test: **p<0,01. (**J**) Kinetics of Mitosox Green fluorescence in H9C2 treated or not with iCT, S3QEL2, or a combination of S3QEL2 and iCT. One-way ANOVA: ns; not significant. (**K**) Histogram of viable H9C2 cell counts, comparing untreated cells to those treated with iCT, S1QEL1, or a combination of iCT and S1QEL1, measured with a Malassez counting chamber (n=6). Brown-Forsythe and Welch ANOVA test: *p<0,05; ***p<0,001. (**L**) Kinetics of Mitosox Green fluorescence before and after pre-treating H9C2 cells with S1QEL1 followed by iCT treatment, or after iCT treatment alone (n=5) One-way ANOVA: *p<0.05.

To precisely determine the electron flow within the ETC responsible for toxic ROS generation following iCT, we used specific inhibitors targeting complexes I and III of the respiratory chain. These included nonlethal doses of complex III inhibitors, such as antimycin A, which is specific to its activity, and S3QEL2 (S3), which specifically inhibits its ROS production. Neither of these two inhibitors could significantly reduce iCT toxicity (Fig. 4I and Fig. S3B). S3 failed to block complex III ROS production (Fig. 4J and S3C). Using rotenone, a well-known complex I inhibitor, we observed restoration of cell viability, accompanied by a reduction in mitochondrial ROS levels (Fig. S3D-F), opening the possibility that RET may play a central role in the observed toxicity (*35*). S1QEL1.1 (S1), an inhibitor specifically targeting ROS production at the IQ site of complex I during RET, completely reversed cell death induced by iCT treatment (Fig. 4K) (*36*). In line with improved viability, the S1 inhibitor significantly reduced iCT induced ROS production (Fig. 4L and S3G).

These experiments provide the first evidence that RET can be induced by chemotherapeutic treatments, leading to pathological ROS production, while highlighting SDH inhibition with malonate as a promising therapeutic solution.

### Malonate treatment prevent iCT-induced cardiotoxicity *in vivo*

To demonstrate the cardioprotective effects of malonate, we implemented *in vivo* a treatment protocol spanning three weeks of injections with the dimethyl version of the SDH inhibitor, encompassing the week of chemotherapy administration. Cardiac function was subsequently analyzed two weeks post-iCT treatment (Fig. 5A). The heart-to-tibia length ratio showed an improvement in the combination treatment group (iCT + DM-Malonate) compared to the iCT-only group, indicating a potential cardioprotective effect, while body weight remained marginally affected (Fig. 5B and Fig. S4A).

**Fig. 5.**
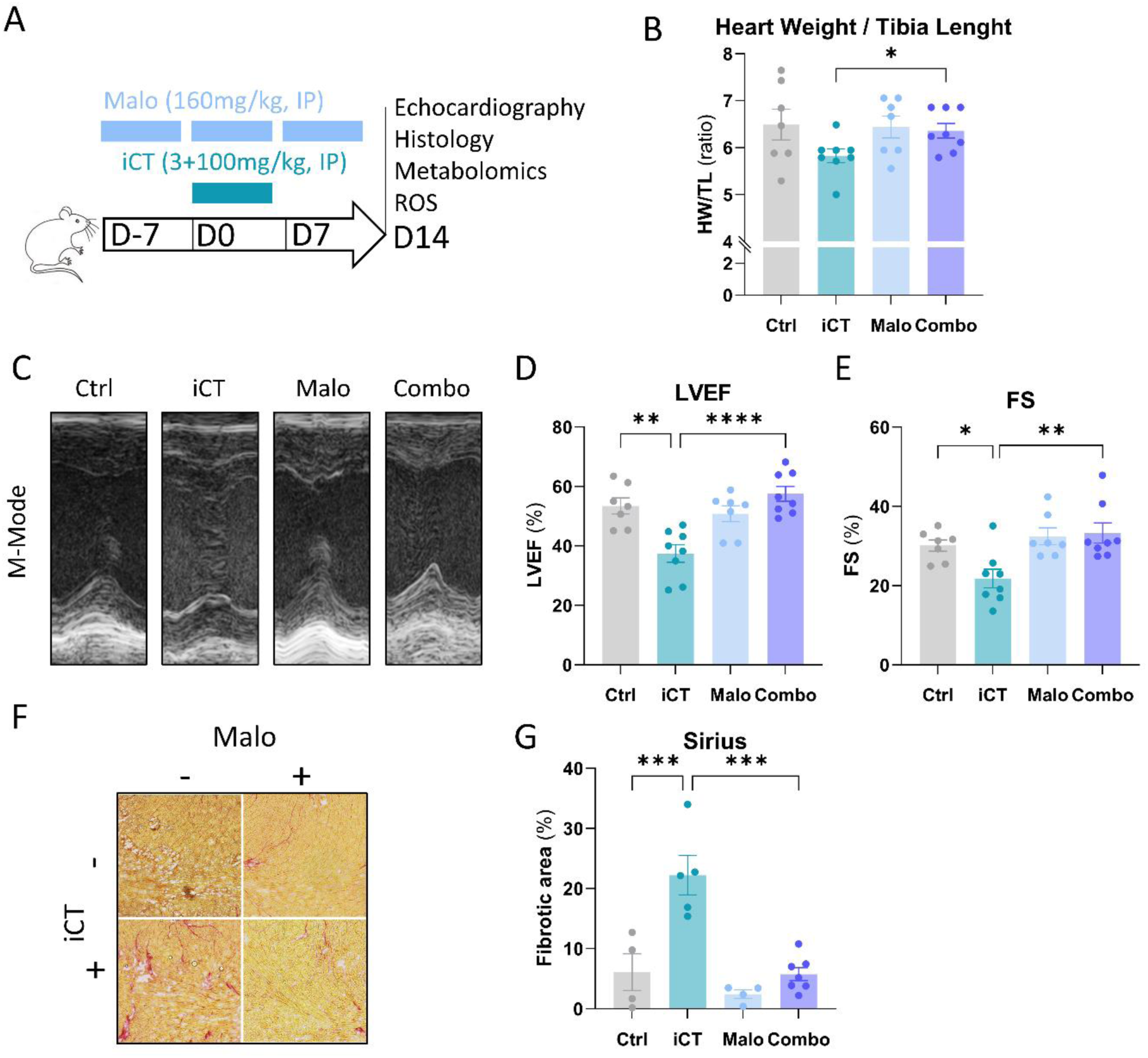
Malonate provides cardioprotection against the harmful effects of Ict. (**A**) Experimental protocol for C57Bl/6 mice: iCT administered for 5 days, with Malonate treatment starting 7 days prior to iCT, continuing throughout iCT treatment, and extending 7 days post-iCT. Echocardiography monitoring was performed on day 14. (**B**) Ratio of heart weight (HW) to tibial length (TL) for untreated mice (n=7) and those treated with iCT (n=8), malonate (n=7), or both iCT and Malonate at day 14 (n=8). One-way ANOVA: *p<0,05. (**C**) Representative M-mode echocardiographic images of mouse hearts, either untreated or treated with iCT, Malo, or the combination of iCT and Malonate. (**D**) Histogram of Left Ventricular Ejection Fraction in untreated mice (n=7) or those treated with iCT (n=8), Malo (n=7), or the combination of iCT and Malo (n=8), measured by echocardiography on day 14. One-way ANOVA: **p<0.01; ****p<0,0001. (**E**) Histogram of fractional shortening in untreated mice (n=7) or those treated with iCT (n=8), Malo (n=7), or the combination of iCT and Malo (n=8), measured by echocardiography on day 14. One-way ANOVA: *p<0.05; **p<0.01. (**F**) Representative images of Sirius Red staining of cardiomyocyte area in heart sections from untreated mice, or mice treated with iCT, Malo, or the combination of iCT and Malo. (**G**) Histogram of quantitative fibrotic area in untreated mice (n=4), or mice treated with iCT (n=5), Malo (n=4), or the combination of iCT and Malo (n=7). One-way ANOVA: ***p<0.001

An assessment of left ventricular systolic function, conducted at equivalent heart rates across groups (Fig. 5C and Fig. S4B), revealed that the significant decrease in ejection fraction and fractional shortening observed in the iCT-induced group was completely restored by the addition of DM-malonate (Fig. 5D and E and Fig. S4C). Similar effects were observed for cardiac output and stroke volume (Fig. S4D and E).

Histological analysis using Sirius Red staining demonstrated a significant increase in cardiac fibrosis following iCT treatment, which was fully reversed with DM-Malonate treatment (Fig. 5F and G).

Metabolomic analysis of heart samples reveals that iCT (as shown in Fig. 2) and DM-malonate induce significant succinate accumulation, whereas their combination does not (Fig. S4F). While succinate accumulation with DM-malonate was expected, the succinate levels observed with the combination therapy suggest that a reversal of SDH activity, in addition to RET, may eliminate excess of electrons by reducing succinate to fumarate. We also observed a near-significant correlation between cardiac succinate concentrations and LVEF across untreated, iCT, and Combo conditions (Fig. S4G). These *in vivo* experiments collectively demonstrate that malonate provides cardioprotection against the toxic effects of the standard chemotherapeutic combination used for AML treatment.

### Malonate enhance therapeutic efficacy of iCT against acute myeloid leukemia

Acute Myeloid Leukemia (AML) blasts heavily rely on mitochondrial oxidative metabolism to withstand treatment, as demonstrated in previous studies (*21, 24, 25, 37*). Therefore, we hypothesized that malonate could be a novel therapeutic strategy to both reduce the risk of cardiac complications and enhance the response to chemotherapy.

Transcriptomic study of viable blasts from MLL/AF9 model treated with the iCT combination (at the same concentrations used in the current study) showed an enrichment of signatures related to mitochondrial metabolism and ROS production (Fig. 6A; GSE139159). Additionally, AML patients from TCGA dataset overexpressing SDHA have a shorter overall survival compared to those with lower SDHA expression (Fig. 6B)(*38*).

**Fig. 6.**
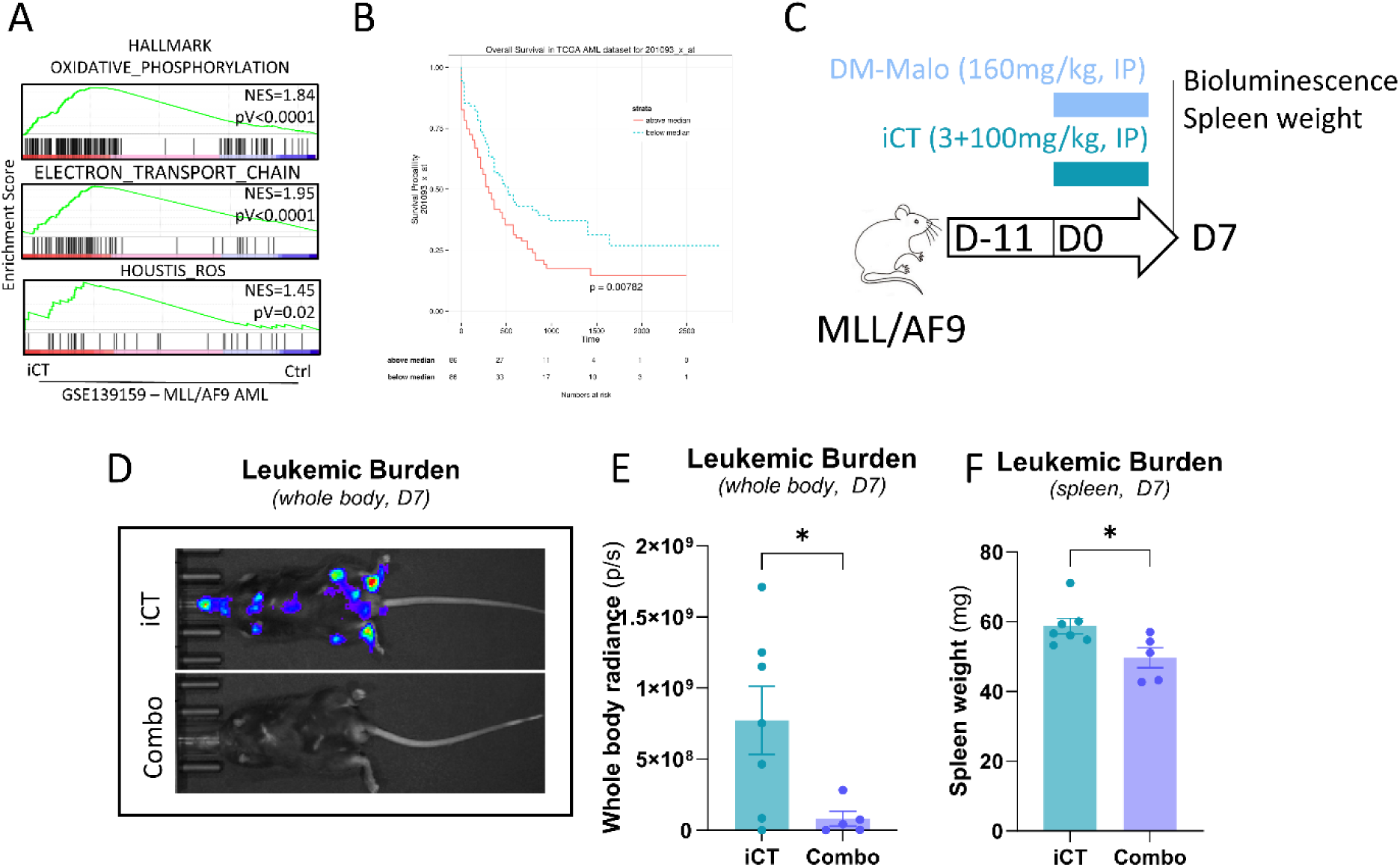
Malonate Enhances iCT-Induced Tumor Burden Reduction in Acute Myeloid Leukemia. (**A**) Gene Set Enrichment Analysis (GSEA) of MLL/AF9 cells treated with iCT. (**B**) Overall survival of AML patients with high SDHA expression compared to those with low SDHA expression. Log-rank test. (**C**) Experimental protocol for C57Bl/6 mice injected with MLL/AF9 cells to develop tumor burden over 11 days. Tumor burden was assessed by bioluminescence before and after 7 days of treatment with iCT alone or in combination with Malonate. (**D**) Representative images of MLL/AF9 cell bioluminescence in mice treated with iCT alone or in combination with Malonate (Combo) (**E**) Histogram of whole-body radiance quantifying tumor burden in mice injected with MLL/AF9 cells and treated with iCT alone (n=7) or in combination with Malonate (n=5). Welch’s t test: *p<0,05 (**F**) Histogram of spleen weight representing leukemic burden in AML mice after treatment with iCT alone (n=7) or in combination with Malonate (n=5). Welch’s t test: *p<0,05.

Next, we used the exact same model from the previous figures (Fig. 6C). However, seven days after treatment with either iCT, malonate, or their combination, we observed an excessive tumor burden for the control and malonate groups necessitating their sacrifice. This clearly highlights the effectiveness of the chemotherapy combination and the lack of antileukemic effect for malonate when used alone. Subsequently, bioluminescence analysis of the leukemic cells revealed residual foci of chemoresistant tumor cells in the iCT-treated mice. In contrast, mice from the combo group showed no detectable or weak signal, demonstrating the potent antileukemic effect of the iCT and malonate combination (Fig. 6D and E). Finally, spleen weights in the combo group were significantly reduced compared to iCT group, reflecting a lower tumor burden in this organ, which serves as a niche for leukemic cells (Fig. 6F).

Thus, the addition of malonate during iCT treatment for AML provides a remarkable cardioprotection against the toxic effect of iCT while reducing tumor burden. Adding malonate to usual chemotherapy is a valuable strategy to enhance the outcomes and benefit-risk balance for AML patients.

## Discussion

This study demonstrates that i) the *in vivo* iCT model used in preclinical AML studies provides a valuable framework for exploring chemotherapy-induced cardiotoxicity, ii) highlights that this toxicity is based on reverse electron transport (RET) in mitochondria and can be reversed by inhibiting SDH with malonate, while iii) simultaneously enhancing the anti-leukemic efficacy of the iCT treatment.

One of the primary limitations in current research on cardiotoxicity relative to chemotherapy is the predominant focus on anthracycline monotherapy like doxorubicin, which often omits to study the complex interactions and metabolic pathways that occur when using combination therapies. Previous studies, have identified oxidative stress and mitochondrial dysfunction as critical mediators of anthracycline-induced cardiac injury (*6, 39, 40*). Recent studies on cardiotoxicity induced by doxorubicin or cytarabine monotherapies, particularly in AML, have shown specific insights on cardioprotective therapies (*28, 41*). As cardiotoxicity occurs mainly during induction with intensive chemotherapy, we replicated this sequence of patient care by using iCT in an *in vivo* experimental model following the 3+7 regimen used for the treatment of AML (*7*). Our results demonstrated that combination chemotherapy was more cardiotoxic than monotherapy, causing a significant reduction in LVEF lasting for at least 15 weeks. While the primary goal of doxorubicin and cytarabine treatments is to induce DNA damage in tumoral proliferating cells, the off-target effects, particularly on mitochondrial metabolism, are distinct, contributing to drug resistance (*42*). Notable mitochondrial effects may include interactions with complex I or mitochondrial iron centers, implication of mitochondrial outer membrane pore, increased OxPhos metabolism or elevated membrane potential driving ROS production (*6, 21, 37*). These effects are particularly toxic to normal cells such as cardiomyocytes.

Through metabolomic analysis, our findings reveal new therapeutic opportunities by demonstrating for the first time that intensive chemotherapy induces the reversion of the electron transport chain (RET). In 2014, Chouchani et al. demonstrated for the first time that malonate could restore cardiac function by inhibiting RET in the context of I/R injury through the inhibition of SDH (*12*). Recent publications have refined the understanding of malonate’s protective mechanisms and optimized its treatment protocol (*14, 43*). Malonate has recently been shown to stimulate the proliferation of adult cardiomyocytes in I/R models; however, this proliferation cannot fully explain the entirety of malonate beneficial effects (*19*). Interestingly, while malonate provided complete cardioprotection when used in combination with doxorubicin and cytarabine, it was ineffective in reversing toxicity when these chemotherapeutic agents were applied as monotherapies. We hypothesize that cytarabine and doxorubicin act synergistically to induce RET, highlighting once again the importance of testing models of combination therapies, as in standard clinical practice. Similarly, this hypothesis may extend to combinations like doxorubicin and cyclophosphamide used in breast cancer treatment. Doxorubicin, known for its electron-exchanging ability with respiratory chain complexes (*44*), may aggravate cardiac dysfunction when cytarabine or cyclophosphamide stimulates OxPhos metabolism (*45*). This synergistic effect on mitochondrial metabolism is illustrated by an increase in succinate, NADH, and ROS production, which are three key hallmarks of the reversal of the electron transport chain that was normalized by RET inhibitors (malonate and S1QEL1.1). Then, it is highly plausible that circulating succinate could serve as an early biomarker of cardiotoxicity, similar to findings in I/R (*46*).

Studies have highlighted the significant role of succinate and its metabolic pathways in various cancers, particularly those with pathogenic variants in succinate dehydrogenase (SDH), such as paragangliomas and pheochromocytomas (*47, 48*). These mutations lead to loss of SDH activity in tumors and to accumulation of succinate, which has been implicated in tumorigenesis through the activation of oncogenic signaling pathways and the stabilization of hypoxia-inducible factors (HIFs) (*49*). Malonate has recently been demonstrated to resensitize resistant cells to BH3 mimetics like venetoclax (*26*) and also proven to be effective against RET with minimal toxic effects in mouse models (*14, 34, 50*). Relative to our results, this suggests that modulating RET in aggressive cancer cells holds promise to curb their proliferation. While RET has rarely been explored in oncology, emerging studies are beginning to reveal its significance. For instance, a 2022 study by Ojha et al. demonstrated that RET plays a role in tumor proliferation in a glioblastoma multiforme model through a Notch-dependent pathway (*51*). Therefore, investigating RET in chemoresistance mechanisms in AML and other cancers offers a new strategy for eliminating residual chemoresistant cells, ultimately enhancing treatment outcomes.

In recent years, it has become increasingly clear that the most resistant cancer cells heavily depend on mitochondrial metabolism for their survival and proliferation (*20, 21, 37*). This understanding led to the development of a potent complex I inhibitor, IACS-010759, which initially showed promise in treating AML (*25*). However, a clinical trial evaluating this inhibitor was ultimately terminated due to brain toxicity and failure to meet primary and secondary endpoints (*52*). This setback, along with prior unsuccessful trials, highlights the challenges of targeting metabolic pathways while avoiding severe side effects (*53*). In our study, SDH inhibitor malonate circumvent these challenges by inhibiting RET-induced toxicity while simultaneously targeting chemoresistant AML blasts metabolic dependency.

This study has several limitations. Among them, we should note that the lethality observed with the 3+7 regimen may stem from overall toxicity affecting organs in addition to the heart. Furthermore, some demonstrations had to be carried out using the imperfect H9C2 rat cardiomyoblast cell line. Despite its limitations, this cell line proved robust as it replicated all unique aspects of reverse electron transport, making it an ideal model for our study. However, the exact molecular pathways leading to RET require further elucidation. The specific contributions of RET and the reversal of SDH activity to iCT-induced toxicity remain unclear, although it is likely that both processes are activated to manage the excess of electrons generated upon chemotherapy treatment. The MLL/AF9 AML model used is particularly aggressive and does not represent all facets of the AML disease. Additionally, the short follow-up period also limits our ability to assess long-term outcomes of SDH inhibition in the context of cancer therapy and cardioprotective strategies. Therefore, in future studies, it would be advantageous to test the malonate-iCT combination on more robust models, such as patient-derived xenografts (PDX) with extended follow-up period. These aspects will be further explored. Additionally, while malonate proved effective in our models, its application on clinical practice needs thorough investigation, particularly regarding its specificity and potential off-target effects.

In conclusion, our study reveals the crucial role of RET in chemotherapy-induced cardiotoxicity, demonstrating therapeutic potential of malonate to mitigate the cardiotoxicity but also to enhance the efficacy of combination therapies. These findings highlight the mitochondrial metabolism as a therapeutic target, especially in the context of combination therapies, that could significantly improve treatment outcomes for patients facing the dual challenges of cancer and chemotherapy-related cardiotoxicity.

## Materials and methods

### Cell culture and drug treatment

H9C2 rat myoblast cells (passage 5 to 15) were cultured in monolayers in MEMα medium (Gibco), supplemented 10% fetal bovine serum (Sigma), 100 U/ml penicillin and 100 μg/ml streptomycin (Sigma) at 37°C in a humidified atmosphere of 5% CO_2_. At 70% confluence, cells were detached with trypsin-EDTA (Thermo Fisher, France), seeded in 24-well plates at 70,000 cells per well, and treated for 24 hours with 0,4 µM Doxorubicin, 4 µM Cytarabine, 100 µM Malonic acid, 5 mM Diethyl or Dimethyl Succinate, 0,1 µM S1QEL1, or 0,1 µM S3QEL2, either alone or in combination.

Primary cardiomyocytes were isolated from C57BL/6 wild-type mice using the Langendorff method, which involved retrograde perfusion of the heart with enzymatic solutions to dissociate the myocardial tissue as described by Liu and colleagues (*54*).

### Animal experiments and *in vivo* treatment

Experiments were conducted on male C57Bl/6 mice aged of 8 to 12 weeks, which were treated with an intravenous injection of Doxorubicin into the tail vein at doses of either 5 mg/kg or 3 mg/kg over 3 days, and an intraperitoneal injection of Cytarabine at doses of either 50 mg/kg or 100 mg/kg over 7 or 5 days respectively. Each experiment included 5 mice per group: untreated, treated with Doxorubicin (Doxo), treated with Cytarabine (AraC), or treated with both Doxorubicin and Cytarabine (iCT). For the diethyl succinate experiment, an intraperitoneal injection of 160 mg/kg was administered daily for 14 days, starting on the same day as the iCT treatment. Each group, including those receiving iCT alone, consisted of 6 mice. For the dimethyl malonate experiment, an intraperitoneal injection of 160 mg/kg was administered 7 days before the iCT treatment, throughout the iCT treatment period and 7 days after. At the end of the treatment, cardiac function was assessed by echocardiography. The animals were then immediately sacrificed using cervical dislocation. After perfusion with PBS, hearts were collected for histological and metabolomic analyses.

Male C57BL/6 mice were injected intravenously via the retroorbital route with luciferase-tagged MLL/AF9 cells to induce acute myeloid leukemia. After 11 days of tumor growth, mice were treated daily with iCT (3 or 100 mg/kg) alone or in combination with Malonate (160 mg/kg) via intraperitoneal injection for 7 days. At the end of the treatment period, tumor burden was assessed by bioluminescence using IVIS imaging system. The mice were sacrificed by cervical dislocation and their hearts and spleens were weighed and collected for further analysis.

### Doppler echocardiography

Cardiac function was evaluated *via* non-invasive echocardiography. Echocardiography was carried out on lightly anaesthetized (1% isoflurane in air) mice placed on a heating pad. The left ventricular dimensions were obtained during Time Movement mode acquisition from the parasternal short-axis view at the midventricular level using a Vevo2100 echograph and a 40 MHz transducer (M550; Fujifilm Visualsonics, Tokyo, Japan). Left ventricular ejection fraction (LVEF) was analyzed in a longitudinal view, and fractional shortening (FS) was assessed using M-mode echocardiography, with all images analyzed offline using VevoLab software (Fujifilm VisualSonics).

### Sirius red staining

The center section of the heart was slowly frozen in OCT (VWR, Radnor; #361603E) using an −80 °C isopropanol bath. Briefly, hearts were transversely sectioned using a cryotome (Thermo NX50; Thermo Fisher Scientific) rinsed quickly in 1 change of acetic acid solution prior to mounting in 95% alcohol with 2 changes in Histoclear solution (Sigma-Aldrich) followed by mounting in synthetic resin. Slides were fixed in 4% PFA in 0.1 M sodium phosphate buffer pH 7.4 for 10 minutes before staining. Picrosirius red (PSR) staining was performed to detect fibrosis according to the protocol of Junqueira et al(*55*). Total fibrotic areas were quantified by using NIH ImageJ software after scanning on a NanoZoomer (Hamamatsu Photonics).

### GSEA (gene set enrichment analysis)

GSEA was performed using the open-source GSEA v.4.0.0 software from the BROAD Institute (http://www.broadinstitute.org/gsea) to identify sets of genes with consistent differences between vehicle-and chemotherapy-treated cells or mice across the datasets detailed in the legends (*56*). This approach evaluates whether predefined gene sets display statistically significant, coordinated changes between two biological states, providing insights into the biology underlying complex datasets.

### Sample preparation

Heart samples weighing approximately 100 mg were homogenized in liquid nitrogen using a mortar and pestle to obtain a fine powder. The resulting powder was transferred to an empty Eppendorf tube, and 450 µL of methanol (pre-chilled at -20°C) was added to the tube. The mixture was then homogenized for 5 minutes at 4°C to extract the metabolites. Following homogenization, the sample was centrifuged at 21,100 g for 10 minutes at 4°C, and the supernatant was carefully collected. The supernatant (400 µL) was diluted with an additional 600 µL of methanol. For biphasic extraction, 0.5 mL of the methanol extract was mixed with 0.5 mL of chloroform and 0.25 mL of water in glass tubes. The samples were incubated at 4°C for 15 minutes with shaking, followed by centrifugation at 2,000 g for 15 minutes at 4°C. After centrifugation, three distinct layers were observed: the upper polar metabolites layer, the middle protein layer, and the bottom lipid layer. The upper layer was carefully transferred to a new Eppendorf tube, and the polar metabolites were dried down in a vacuum concentrator (SpeedVac, Thermo Fisher Scientific) and resuspended in 90% acetonitrile. Prior to analysis, the samples were filtered through a 0.2 μm PTFE membrane filter before injection.

### LC-MS and data processing

Liquid chromatography was performed with a Vanquish Neo UHPLC (Thermo Scientific) using ZIC-HILIC trap column (0.3 × 10 mm, 5 μm, Merck) and separation column (0.3 × 150 mm, 3.5 μm, SeQuant Merck). 5 μl of sample or blank was injected onto the trap column 30 seconds before gradient elution. Mobile phase A of the HILIC gradient was 5% acetonitrile, 95% water containing 5 mM ammonium acetate; mobile phase B was 90% acetonitrile, 10% water. The separation proceeded as follows: from 0 to 12.2 min mobile phase B was kept at 100% to elute nonpolar compounds; from 12.2 min to 42.2 min, B was ramped linearly from 100% to 42.5% then decreased to 10% at 44.2 min; B was held at 10% from 44.2 min to 49.2 min to remove salt from the column; we kept B at 100% from 49.2 min to 64 min to regenerate the water layer of the column. The solvent flow rate was set at 3 μl/min. The column temperature was maintained at 40°C.

Mass spectrometry data were acquired using Orbitrap Exploris 240 (Thermo Scientific) in full scan MS mode without polarity switching. Each polarity was acquired at a resolution of 60,000 (at m/z = 200); default settings were used for the automatic gain control target and maximum injection time; microscans was set to 2 and the scan range to 70–800 Daltons. MS source parameters were as follows: capillary voltage (V): 3400 for positive ion mode and 2300 for negative ion mode; sheath gas (Arb/ arbitrary unit): 3; auxiliary gas (Arb): 2; sweep gas (Arb): 0; ion transfer tube temperature: 320 °C.

Raw data in .raw format was converted to .mzXML format using ProteoWizard (DOI: 10.1038/nbt.2377). The .mzXML files were imported and processed in MZmine 2.53 (DOI:10.1038/s41587-023-01690-2). Metabolite detection was performed in targeted feature detection mode by searching exact masses of known metabolites. The parameters for feature extraction were as follows: mass detection (exact mass, noise 1.0E3); targeted feature detection (intensity tolerance 80, noise 1.0E3, m/z tolerance 2 ppm); smoothing (filter width 11). The feature lists were then manually inspected to assess peak shape and peak integration before being exported for data analysis.

### Mitochondrial ROS quantification

Untreated H9C2 cells were seeded into 96-well plates at a density of 20,000 cells per well and incubated for 24 hours. Following incubation, cells were washed twice with PBS and then treated with MitoSOX Green for 20 minutes. MitoSOX Green is a fluorogenic probe that emits fluorescence upon oxidation by superoxide, a reactive oxygen species (ROS) in the mitochondria. To measure mitochondrial ROS production, fluorescence intensity was monitored using the Incucyte SX G/O/NIR Optical Module (Sartorius) with a 20X objective. Baseline fluorescence was recorded before any treatment. Subsequently, fluorescence was measured in the same wells immediately after the first injection of 100 µM Malonate, 1 µM S1QEL1, 1 µM S3QEL3, or 1 µM Rotenone. Additional measurements were taken following a second injection of iCT in the same wells to assess changes in ROS production over time. For histograms, quantification was performed by calculating the mean of each phase.

### Succinate assay

Succinate concentrations in H9C2 cells were quantified using the Succinate Colorimetric Assay Kit according to the manufacturer’s protocol (Sigma-Aldrich, MAK184). The colorimetric measurements were taken using a TECAN 500 microplate reader.

### NAD/NADH Assay

NAD and NADH concentrations in H9C2 cells were quantified using the NAD/NADH Assay Kit according to the manufacturer’s protocol (Sigma-Aldrich, MAK468). Optical density was measured using a BMG Labtech Spectrostar NanoDosage.

### Cell viability assay by Malassez counting

Cell viability was assessed using a Malassez counting chamber. Cells were stained with Trypan blue to distinguish between viable and non-viable cells, and only viable cells were counted.

### Seahorse Extracellular Flux (XF) Cell Mito Stress

Mitochondrial function was evaluated using Seahorse XFe24 Flux Analyzer (Agilent, Santa Clara, CA, USA) by measuring oxygen consumption rate (OCR) and extracellular acidification rate (ECAR). H9C2 cells were seeded on the Seahorse microplate at a density of 5 × 10^4^ cells per well in DMEM supplemented with L-glutamine (2 mM), sodium pyruvate (1 mM) and glucose (10 mM), pH 7.4 for the Mitostress Test. To further define the role of mitochondrial respiration in cell metabolism, the compounds oligomycin (1 μM), carbonyl cyanide 4-(trifluoromethoxy) phenylhydrazone (FCCP, 2 μM) and a mixture of rotenone and antimycin A (RAA, 1 μM) were serially injected to measure basal, maximal OCR and ATP production. All reagents came from Sigma-Aldrich (Saint-Louis, MI, USA).

### Mitochondrial membrane potential analysis by flow cytometry

Functional status of mitochondria was evaluated by MitoTracker Red CMXRos dye (Thermo Fisher Scientific, Waltham, MA, USA, Cat. No. M7512), depending on mitochondrial membrane potential of living cells. H9C2 cells were seeded in a 6 well plate at a density of 300.000 cells per well and incubated overnight at 37°C with 5% CO_2_. The following day, cells were detached using trypsin-EDTA (Thermo Fisher, France) and then incubated in a cytometry tube with 1 mL of MitoTracker Red at a concentration of 1:500,000 for 30 minutes at 37°C with 5% CO_2_. After incubation, cells were washed twice and resuspended in 1 mL of warm HBSS. The mitochondrial membrane potential was initially evaluated by flow cytometry (Cytek Aurora CS) in untreated H9C2 cells. Doxorubicin fluorescence was excluded using the spectral acquisition capabilities of this specific cytometer.

### Western blotting

For Western blot analysis on mice tissue, proteins from frozen heart ventricular apex were extracted using a FastPrep-24 instrument (MP Biomedical) following the manufacturer’s recommendations. After quantification with DC protein assay (Bio-Rad), proteins were run on Mini protean TGX STAINFREE 4%-15% gel (Bio-Rad) and transferred to Amersham Protran 0.2µM nitrocellulose blotting membrane (10600004, GE Healthcare) with Mini Trans-Blot transfer system. Membranes were blocked for 1 hour at room temperature with 10% albumin in PBS containing 0.05% Tween 20 (Sigma-Aldrich) and then incubated with the appropriate primary antibody overnight at 4 °C. Primary antibody used here: SDHA at 1:1000 dilution from Cell Signaling (#11998). After PBS containing 0.05% Tween 20 (Sigma-Aldrich) washes, membranes were incubated with the appropriate HRP-conjugated secondary antibody for 1 hour at room temperature. Blots were quantified on a ChemiDoc imaging system (Bio-Rad) using the previously described protocol (*57*) and normalized to actin expression.

### Statistical Analysis

Statistical analyses among different groups were performed with GraphPad Prism 10. Comparisons between two groups were made using Mann-Whitney Test while multiple group comparisons were carried out by Brown-Forsythe and Welch ANOVA test, One-way ANOVA or Two-way ANOVA analysis. Statistical analysis of the normalized data was performed using a one-sample t-test. Data are presented as mean ± SEM; ∗p < 0.05; ∗∗p < 0.01, ∗∗∗p < 0.001; ∗∗∗∗p < 0.0001.

## Acknowledgments

We thank all the members the facilities of the “Institut des maladies métaboliques et cardiovasculaire” and from the Deduve Institute who contributed to this work. The authors acknowledge members of the UMS 006 CREFRE-Rangueil experimental zootechny for animal housing and CREFRE-ENI for giving us the access to the Vevo2100; UMS 006 Inserm/UT3/ENVT, Anexplo/Genotoul Plateform. We also thank INSERM, University of Toulouse and CHU Toulouse for their financial support.

## Author contributions

Conceptualization: CP, FS, TF

Methodology: CP, NVG, FS, TF

Investigation: CP, YL, SB, YSM, LL, DM, TF

Visualization: CP, FS, TF

Funding acquisition: FS, TF

Project administration: FS, TF

Supervision: FS, TF

Writing – original draft: CP, FS, TF

## Data and materials availability

All data associated with this study are present in the paper or supplementary Materials.

