## Supplemental figures for "Intensive Chemotherapy Induces Cardiotoxicity via Reverse Electron Transport"

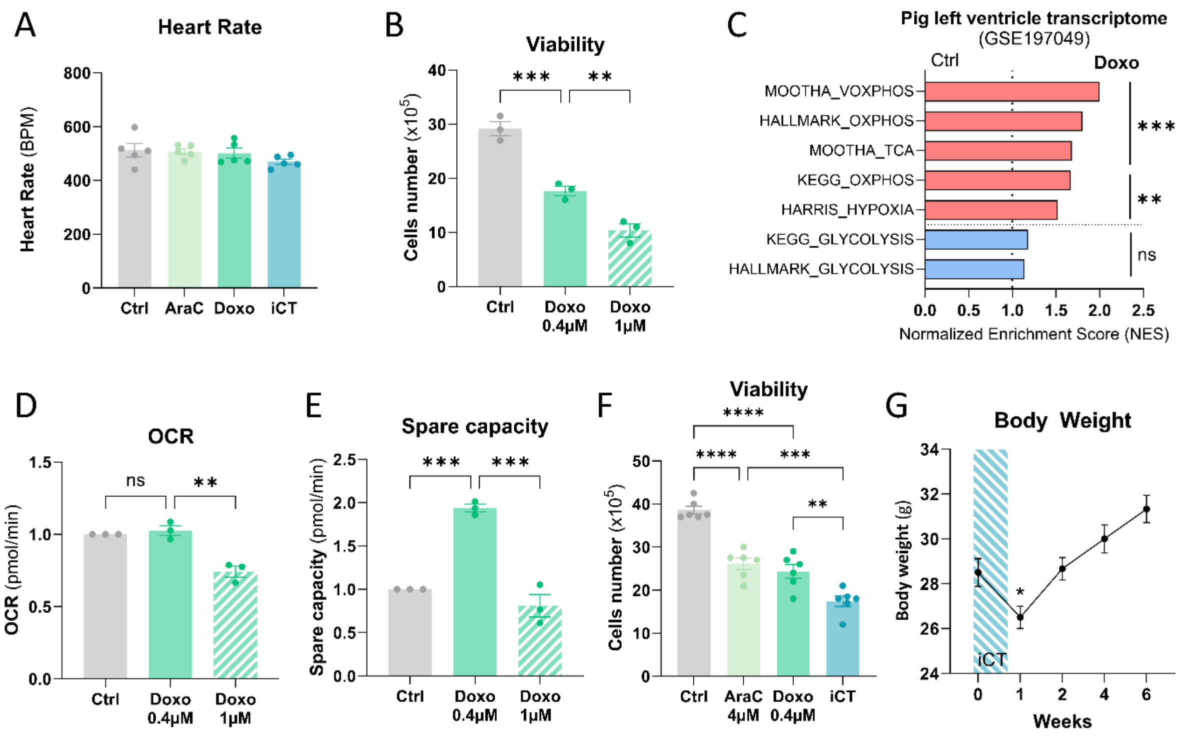


**Fig. S1**

**Fig. S1. Doxorubicin and cytarabine combination drive major cardiotoxic effects**

(**A**) Mouse heart rate after 7 days of treatment. (**B**) Histogram of viable cell counts in untreated H9C2 cells versus those treated with Doxo at 0.4 or 1µM, measured using a Malassez counting chamber (n=3). One-way ANOVA: **p<0.01; ***p<0.001. (**C**) Transcriptomic analysis of doxorubicin-treated pig left ventricule (GSE197049) (**C**) Oxygen Consumption Rate (OCR) measured by Seahorse MitoStress Test in Doxorubicin-treated versus untreated H9C2 Cells (n=3). One-way ANOVA: ns: not significant; **p<0.01. (**E**) Spare capacity measured by the Seahorse MitoStress Test in Doxorubicin-treated versus untreated H9C2 cells (n=3). One-way ANOVA: ***p<0.001. (**F**) Histogram of Viable Cell Counts in untreated versus AraC, Doxo, or iCT treated H9C2 cells measured using a Malassez counting chamber (n=6). One-way ANOVA: **p<0.01; ***p<0.001; ****p<0.0001. (**G**) Kinetics of Mouse Body Weight Over 6 Weeks Following AraC, Doxo, and iCT Treatments as Per the 5+3 Protocol (n=6 mice per group). One-way ANOVA, *p<0,05; **p<0,01.

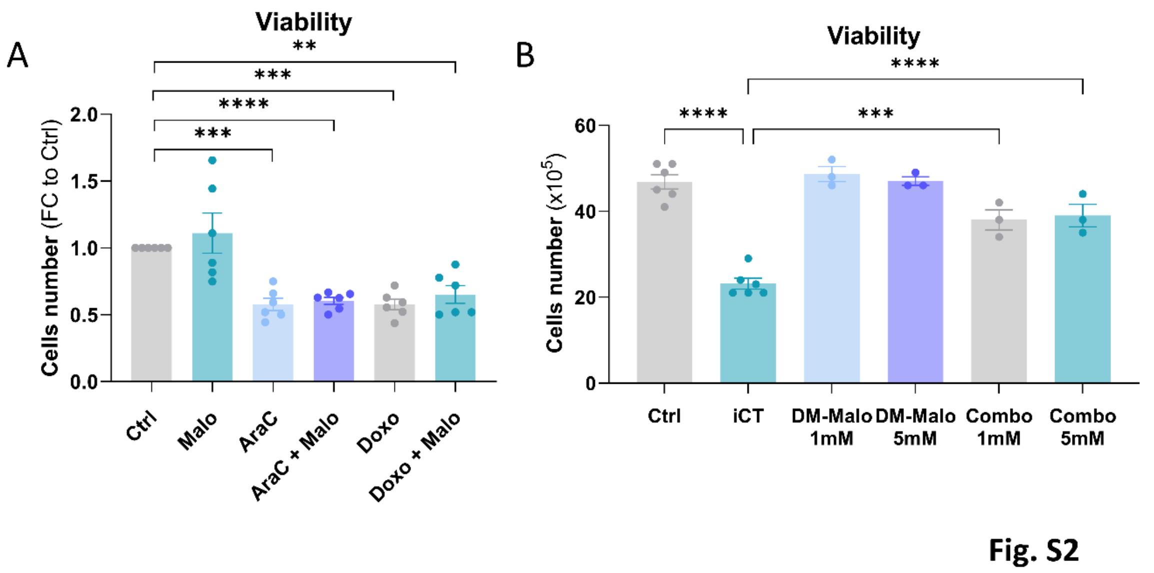


**Fig. S2. Malonate reverses iCT toxicity induced by succinate accumulation**

(**A**) Histogram of viable cell counts in untreated versus Malo, AraC, Doxo, AraC + Malo and Doxo + Malo-treated H9C2 cells, measured using a Malassez counting chamber (n=6). One-sample t-test: **p<0.01; ***p<0.001; ****p<0.0001. (**B**) Histogram of viable cell counts in untreated versus iCT, Dimethyl malonate (DM-Malo) or the combination of iCT and DM-Malo treated H9C2 cells, measured using a Malassez counting chamber (n=3). One way ANOVA: ***p<0.001; ****p<0.0001. (**C**)

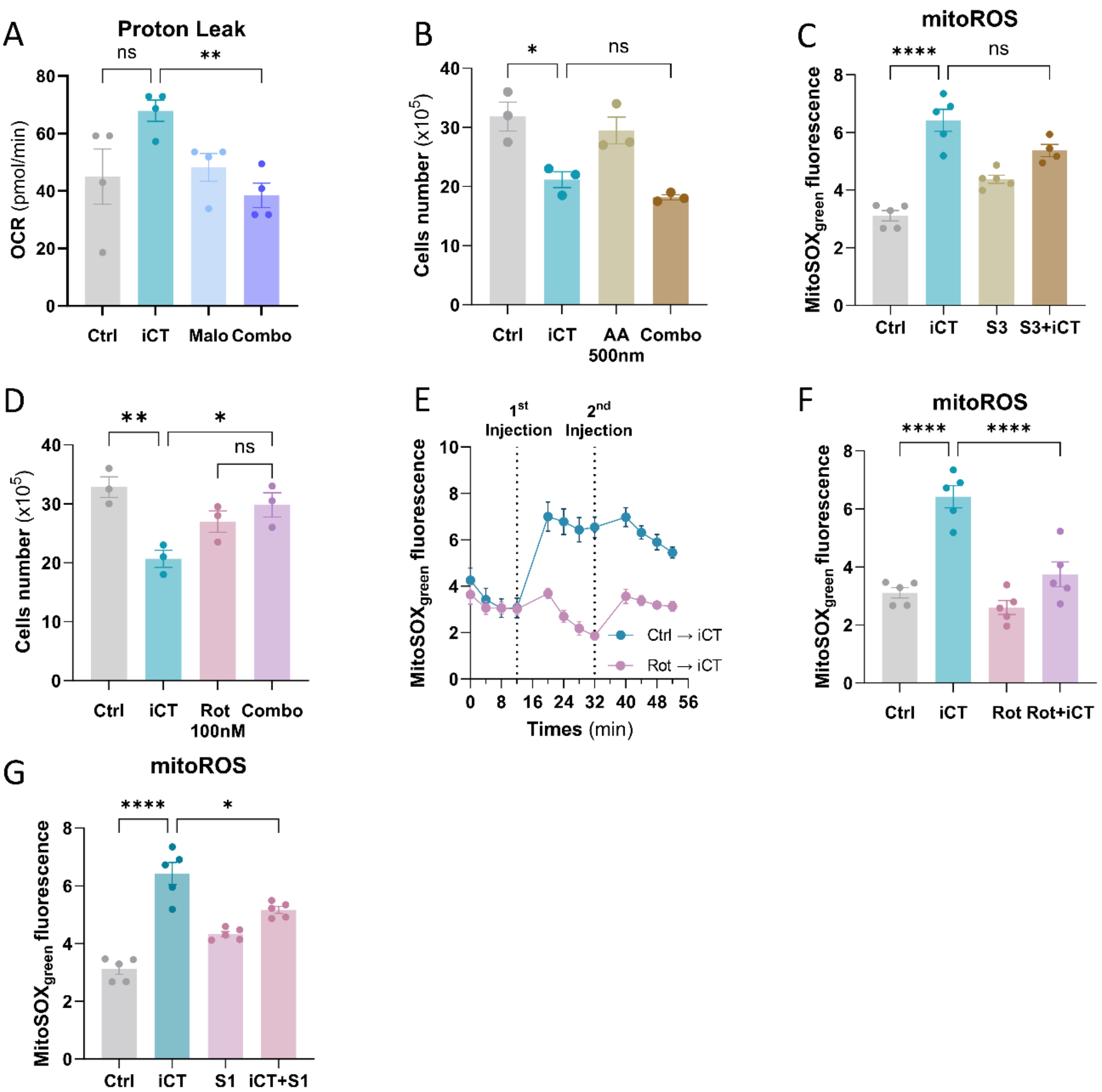


**Fig. S3**

**Fig. S3. iCT increases ROS production through mitochondrial reverse electron transport**

(**A**) Proton leak histogram measured by Seahorse MitoStress Test in H9C2 cells either untreated or treated with iCT, malonate, or the combination of iCT and malonate (Combo) for 24 hours (n=4). Brown-Forsythe and Welch ANOVA test, *p<0,05. (**B**) Histogram of viable cell counts in untreated versus iCT, Antimycin A (AA) or the combination of iCT and AA treated H9C2 cells, measured using a Malassez counting chamber (n=3). One way ANOVA: ns: not significant; *p<0.05. (**C**) Histogram quantification of MitoSOX Green fluorescence representing mitochondrial ROS before and after pre-treating H9C2 cells with S3QEL2, followed by iCT treatment, or after iCT treatment alone (n=5). One-way ANOVA: ****p<0.0001; ns : not significant. (**D**) Histogram of viable cell counts in untreated versus iCT, Rotenone (Rot) or the combination of iCT and Rot treated H9C2 cells, measured using a Malassez counting chamber (n=3). One way ANOVA: *p<0.05; **p<0.01. (**E**) Kinetic (**F**) Histogram quantification of MitoSOX Green fluorescence representing mitochondrial ROS before and after pre-treating H9C2 cells with rotenone, followed by iCT treatment, or after iCT treatment alone (n=5). One-way ANOVA: *p<0.05; **p<0.01. (**G**) Histogram quantification of MitoSOX Green fluorescence representing mitochondrial ROS before and after pre-treating H9C2 cells with S1QEL1.1, followed by iCT treatment, or after iCT treatment alone (n=5). One-way ANOVA: *p<0.05; ****p<0.0001.

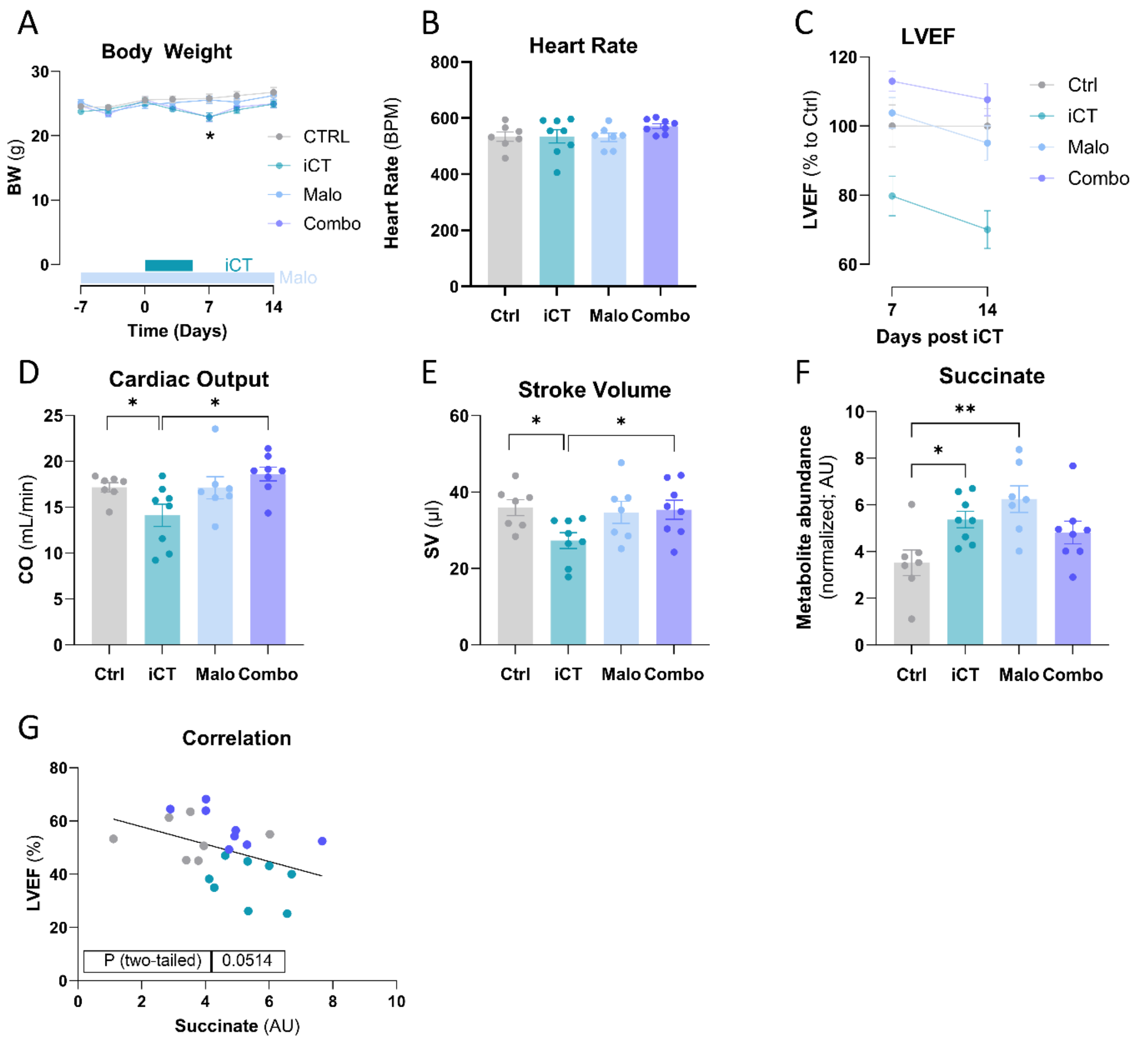


**Fig. S4**

**Fig. S4. Cardioprotective effects of malonate in mitigating iCT-induced cardiotoxicity *in vivo***

(**A**) Body weight monitoring over 21 days in untreated mice (n=7) and those treated with iCT (n=8), malonate (n=7), or the combination of iCT and malonate (n=8). One-way ANOVA: *p<0.05. (**B**) Heart rate measurement by Doppler echocardiography in untreated mice (n=7) and those treated with iCT (n=8), malonate (n=7), or the combination of iCT and malonate. One-way ANOVA. (**C**) Echocardiographic Monitoring of Left Ventricular Ejection Fraction in untreated mice (n=7) or those treated with iCT (n=8), malonate (n=7), or the combination of iCT and malonate (n=8) at 7- and 14-days post iCT. (**D**) Cardiac Output and (**E**) Stroke Volume histogram of untreated mice (n=7) and those treated with iCT (n=8), malonate (n=7), or the combination of iCT and malonate (n=8) measured by echocardiography after 21 days of treatment. One way ANOVA and Welch's t test: *p<0.05. (**F**) Histogram of succinate metabolite abundance in iCT (n=8), malonate (n=7) and combo-treated mouse hearts (n=8) compared to controls (n=7). One-way ANOVA: *p<0.05; **p<0,01. (**G**) Correlation between LVEF and succinate concentration in Control (n=7), iCT (n=8) and combo-treated mouse hearts (n=8). Pearson: p=0.0514.
